# Assessing the socio-economic value of sea turtles to the Maldives’ tourism industry in 2022 (post-pandemic)

**DOI:** 10.64898/2026.04.21.720026

**Authors:** Julian Gervolino, Olivia Forster, Isha Afeef, Risha Ali Rasheed, Abdulla Hameed, Ibrahim Inan, Dan Nashid, Claire Petros, Jane R. Lloyd, Stephanie Köhnk

## Abstract

The non-consumptive use of sea turtles has become a rapidly growing sector of the global tourism industry and is increasingly recognised as an important source of economic benefit for coastal communities. In the Maldives, however, the socio-economic value of sea turtles remains poorly quantified. Building on a preliminary survey conducted in 2019, this study provides a comprehensive assessment of the socio-economic value of sea turtles to the tourism industry, post-pandemic, while identifying sea turtle viewing hotspots with high tourism pressure that may require stricter regulation. Questionnaires collected information on operations related to sea turtle excursions in 2022, including the estimated direct spend on sea turtle excursions and the perceived value of sea turtles to tourists and operators. Results include responses from 117 tour operators across 20 atolls in the Maldives, with 68% from resort operators, 27% from local island operators, and 5% from liveaboards. Maldivian nationals made up 55% of respondents and 78.8% of the people working, directly or indirectly, on sea turtle excursions in 2022. The direct revenue generated by sea turtle tourism in the Maldives is estimated at US$10.9–14.4 million in 2022, with individual turtles at high-use sites generating up to US$29,709 per year. These findings underscore the significant economic and social importance of sea turtles to the tourism industry in the Maldives, matching that of manta rays and sharks, and highlights the need for improved conservation efforts to safeguard local sea turtle populations and their associated benefits for Maldivian communities.

## Introduction

The exploitation of sea turtles for subsistence and commercial uses has been well-documented across tropical and subtropical coastal communities for millennia [1–3], and has played a major role in the historic decline of wild sea turtle populations globally [4–6]. In more recent decades, additional anthropogenic pressure from marine pollution [7], industrial fishing [8, 9], habitat loss [10], and climate change [11, 12], has further accelerated these population declines. As a result, six of seven species of sea turtles are currently threatened with extinction [13]. However, conservation efforts have helped slow this decline and, in some regions, contributed to the recovery of certain sea turtle species [14, 15]. In particular, the non-consumptive use of sea turtles in ecotourism is rapidly emerging as an effective conservation strategy, offering both economic benefits for many coastal communities and strong incentives for the protection of these species [16–19].

Over the past decade, wildlife tourism has emerged as a major sector in the global tourism industry, accounting for over 4% of estimated direct global Tourism GDP - approximately five times the value of the illegal wildlife trade [20, 21]. In the Maldives alone, tourism has become the dominant economic sector since its boom in 1972 [22], generating $4.5 billion in revenue from 1.68 million visitors in 2022 [23]. A major attraction for visitors is the chance to experience local marine wildlife through in-water activities such as snorkeling and scuba diving [24]. Sea turtles in particular are a key example of marine wildlife that adds value to the experience of visitors to the Maldives, either through in-water encounters or witnessing nesting and hatching events [25]. Despite their apparent value for tourism, all five of the seven species found in the Maldives are listed on the International Union for Conservation of Nature (IUCN) Red List of Threatened Species [26–30].

Sea turtles, their nests, and their habitats have been legally protected in the Maldives since 2016 under the Environmental Protection and Preservation Act (Number 4/93 Section A - Conservation of Biodiversity). The two most commonly found species, hawksbill turtles (*Eretmochelys imbricata*) and green turtles (*Chelonia mydas*), likely have the highest interactions with tourists as they are regularly encountered in reefs, lagoons, and seagrass meadows across the country [31]. Despite their apparent abundance, green and hawksbill turtles are listed as ‘Endangered’ and ‘Critically Endangered’ respectively on the Maldives National Red List of Threatened Species [32, 33]. The major threats facing these species in the country include the loss of nesting and foraging habitat, entanglement in marine debris, the pet trade, and illegal take for egg and meat consumption [34, 35]. Until its ban in 1996, the trade of hawksbills and hawksbill shells as souvenirs was also prominent in the country [31, 36].

Non-consumptive wildlife tourism can have major implications for sea turtle conservation [37–39]. By providing alternative sources of income for local communities, sea turtle ecotourism can support livelihoods while discouraging harmful activities that threaten struggling sea turtle populations [16, 40]. While many successful sea turtle ecotourism initiatives have primarily focused on nesting and hatching observations [16, 41, 42], in-water encounters through diving and snorkeling have also been shown to be a major attraction for tourists [43–46].

Existing studies have primarily focused on the socio-economic value associated with sea turtle nesting ecotourism [17, 39, 40, 42, 46, 47] and/or have incorporated Willingness-to-pay (WTP) assessments in their valuation estimates [16, 18, 48]. However, few studies have assessed the direct economic value of sea turtle tourism through in-water activities such as snorkeling and scuba diving [49], both of which are a central part of the Maldives tourism industry [24] and an increasingly popular way of viewing marine wildlife [50].

Within the Maldives, previous research has estimated the annual direct revenue generated from manta ray and shark tourism to be US$8.1 million and US$14.1 million, respectively [51, 52]. However, the first assessment of the economic value of sea turtles in the Maldives was only recently conducted and estimated it to be valued at US$1.08 million in 2019, prior to the disruption from the COVID-19 pandemic [53]. A previous study by Waheed [54] also noted that visitors to the Maldives were willing to pay a surcharge of US$10.50 for a guaranteed sighting of a sea turtle on in-water excursions. These studies highlight the importance of sea turtles to the country’s tourism industry and the potential for sea turtle ecotourism to grow into a major contributor to the Maldivian economy, comparable to that of other marine megafauna. Nevertheless, due to a small sample size (n=21), the 2019 estimate is not representative of the true economic value of sea turtles in the Maldives. This study therefore aims to build upon previous research to provide a more comprehensive assessment of the socio-economic value of sea turtles in the Maldives, post-COVID-19 pandemic. These findings provide empirical evidence of the significant socio-economic value of sea turtles to the Maldivian tourism industry and highlight opportunities to further develop this ecotourism sector in ways that support sustainable economic development and strengthen sea turtle protection and management plans.

### Methodology

#### Data Collection

Social-survey questionnaires were conducted over six months between 1^st^ June 2023 and 30^th^ November 2023. The questionnaires (S1 Appendix) were completed either online, in-person and/or over the phone depending on the preference and availability of the participant. The questionnaire was available in both English and Dhivehi (local language) to enhance accessibility and equitable participation among Maldivian respondents. The aim was to collect survey responses from a representative number of tour operators from each of the 20 administrative atolls in the Maldives. The questionnaires were facilitated by Olive Ridley Project (ORP) staff in the Maldives.

The study was carried out under permit number EPA/2023/PSR-T04 granted by the Environmental Regulatory Authority (formerly Environmental Protection Agency Maldives), and followed the ethical principles of informed consent, confidentiality, and voluntary participation as outlined in the Declaration of Helsinki. The research involved minimal risk and was non-interventional. All participants were contacted by phone prior to completing the questionnaire to be informed of the aim and scope of the study and to provide informed verbal consent to participate. Consent was obtained verbally from respondents rather than in writing prior to administering the questionnaire as to not discourage participation and to streamline the process when collecting high volumes of survey responses. Participation was optional and respondents could decline to answer any questions or withdraw from the survey at any time. Prior to conducting or sharing the questionnaire, participants were informed that their responses would remain confidential and anonymous, and would only be used for research purposes. No personal identifying information is reported in this study.

#### Questionnaire Design

The survey aimed to measure four key indicators that can provide the socio-economic value of sea turtles: i) the perceived value of sea turtles to tour operators, ii) the direct spend on sea turtle trips, iii) the estimated tourism value of a single turtle, and finally iv) perceived importance to tourists compared to other locally found marine megafauna.

To obtain information from operators, a questionnaire was sent out to collect data on their operations during 2022, after the COVID-19 pandemic, which resulted in major restrictions for international tourism and a decline in Maldives’ GDP by 33.5% [55]. The surveys were advertised on social media platforms such as Facebook and Instagram, and were also directed to known tour operators, such as those working at the same resorts as ORP team members, and other acquaintances. The survey was split into four sections to collect information on: i) the operator itself and details of their employees and guests served in 2022; ii) the direct revenue generated by advertised sea turtle dive excursions and iii) advertised sea turtle snorkel excursions (measured in price ranges e.g. $51-75, $76-100 etc.); and iv) how sea turtles featured in their operations in 2022 (e.g. marketing, merchandise, artwork).

#### Data Analysis

The total direct revenue generated from sea turtle tourism in the Maldives in 2022 was estimated only from advertised sea turtle snorkel and dive trips. Revenue from general snorkel or dive trips, or those specifically targeted at seeing other marine megafauna such as sharks or manta rays were not included:

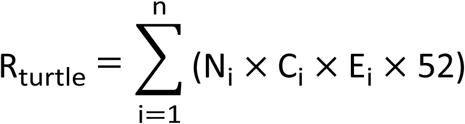

For each tour operator (*n*), the average number of guests per excursion (*N_i_*), average cost per person (*C_i_*), and the average number of excursions per week (*E_i_*) were used to estimate weekly direct revenue from sea turtle excursions. This value was then extrapolated to 52 weeks to obtain an annual estimate for each operator in 2022. The annual estimates across all operators that offered advertised sea turtle excursions were summed to calculate the total direct revenue generated from sea turtle tourism that year (*R_turtle_*). Anderson et al. [51] uses a similar method to estimate annual direct revenue from manta ray tourism in the Maldives. Averages were requested instead of exact figures to facilitate the ease of response and to avoid requiring operators to disclose potentially confidential data, which might have discouraged participation in the questionnaire. As such, we assume that the respondents accounted for seasonal variations in tourism activity when providing these averages. The revenue from any operators that offered advertised sea turtle dive or snorkel or both excursions, but did not complete the pricing and operation questions, were estimated by extrapolating averages of ‘Average trips per week’, ‘Price per guest’ and ‘Guests per trip’ from complete survey responses.

The direct revenue generated by a single turtle in 2022 was estimated for the most popular sea turtle snorkel and dive sites (≥ 3 operator mentions as a sea turtle viewing site). Following a similar approach to Anderson et al. [51], at each site, the number of individuals accounting for 75% of recorded sightings in 2022 was divided by 75% of the total estimated direct revenue generated from all operators frequenting that site. This aims to capture the value of resident sea turtles at a given site while minimizing the influence of incidental sightings of migratory individuals. Sea turtle sightings are recorded through ORP’s photo identification (Photo-ID) research program, in which individual turtles are photographed and identified using their unique facial scute patterns [56]. This survey method does make several assumptions, including: i) all sites listed by each operator (if more than one) are visited equally, ii) operators visit a maximum of three different sites for advertised sea turtle trips, iii) 75% of recorded sightings at a given site are only from resident individuals.

This study does not include the contribution of associated tourism expenditures, such as accommodation, food and other purchases attributed to sea turtle excursions, to the value of sea turtle tourism. The tourism value of sea turtle nesting and hatching experiences are also not included in this study as they are typically offered as free, opportunistic events for tourists in the Maldives to witness, and thus, would be difficult to assess without a WTP survey, as demonstrated by Tisdell and Wilson [57].

## Results

### Survey return

117 tour operators across 20 administrative atolls in the Maldives were surveyed, with about 68% of total responses submitted from resort operators (n = 79), 27% from local island operators (n = 32), and 5% from liveaboards (n = 6). Resort operators had the highest response rates with approximately 47% of resorts surveyed responding to the questionnaire, while local island operators had significantly lower response rates. Only four of the 20 atolls surveyed had no responses, including Haa Alifu, Faafu, Gaafu Dhaalu, and Seenu.

Kaafu atoll (including North and South Malé) made up the largest proportion of survey responses at 29.9% (n = 35), followed by Lhaviyani (11.1%; n = 13), Baa (10.3%; n = 12), and Alifu Dhaalu (10.3%; n = 12), which together accounted for over 60% of responses (Fig 1). 20 different nationalities were recorded in the survey responses, with Maldivians making up 55% of respondents (n = 64) and a large proportion of international respondents from countries such as Italy, France, Germany and the UK.

**Figure 1.**
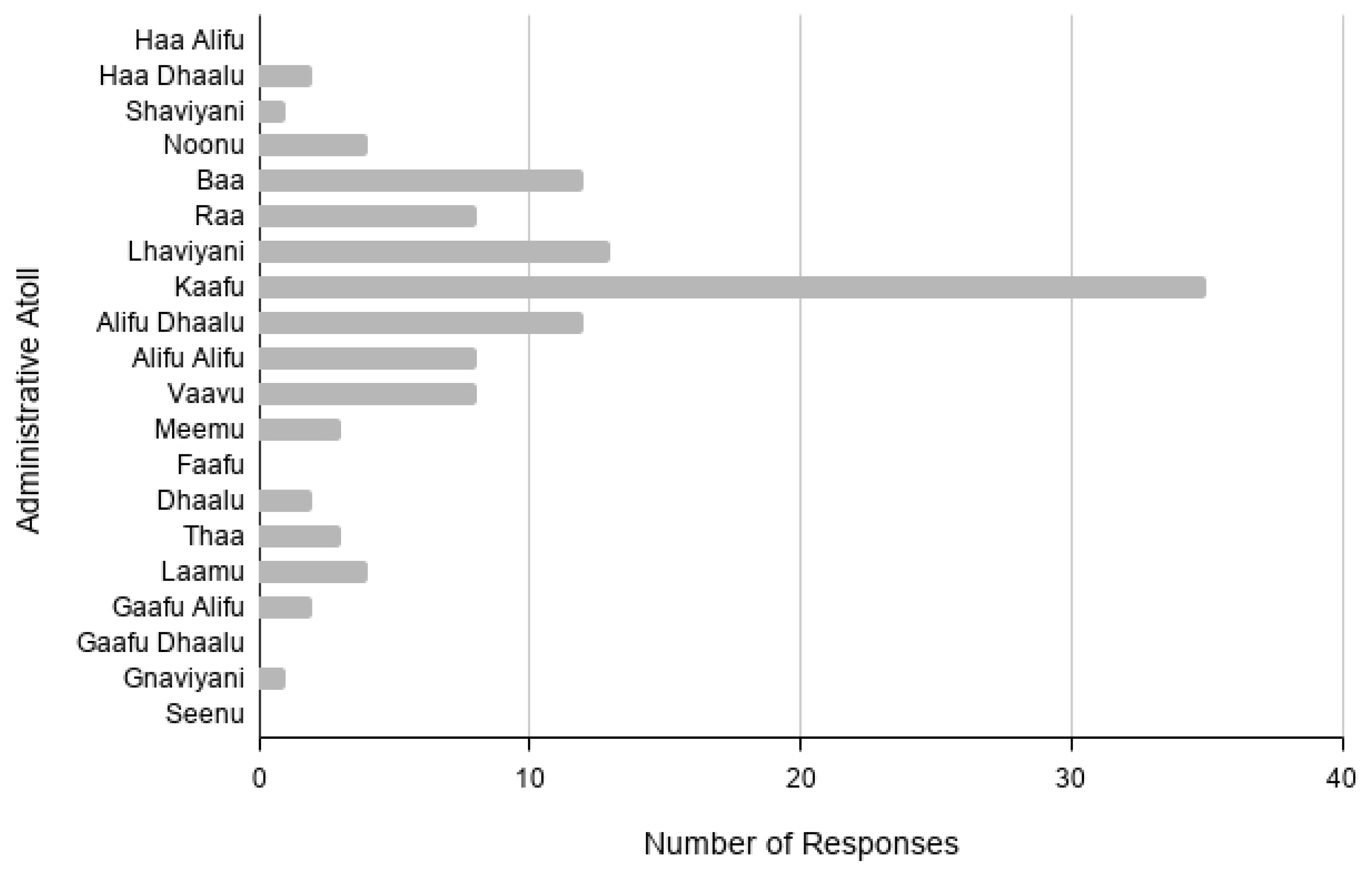
Distribution of survey responses across Maldivian administrative atolls, ordered geographically from north (top) to south (bottom).

Snorkel trips were the most commonly advertised sea turtle excursion among operators in 2022 (n = 51) with significantly fewer operators offering only advertised sea turtle dives (n = 8; Fig 2). 33 operators offered both advertised sea turtle snorkels and dives and 25 operators offered neither.

**Figure 2.**
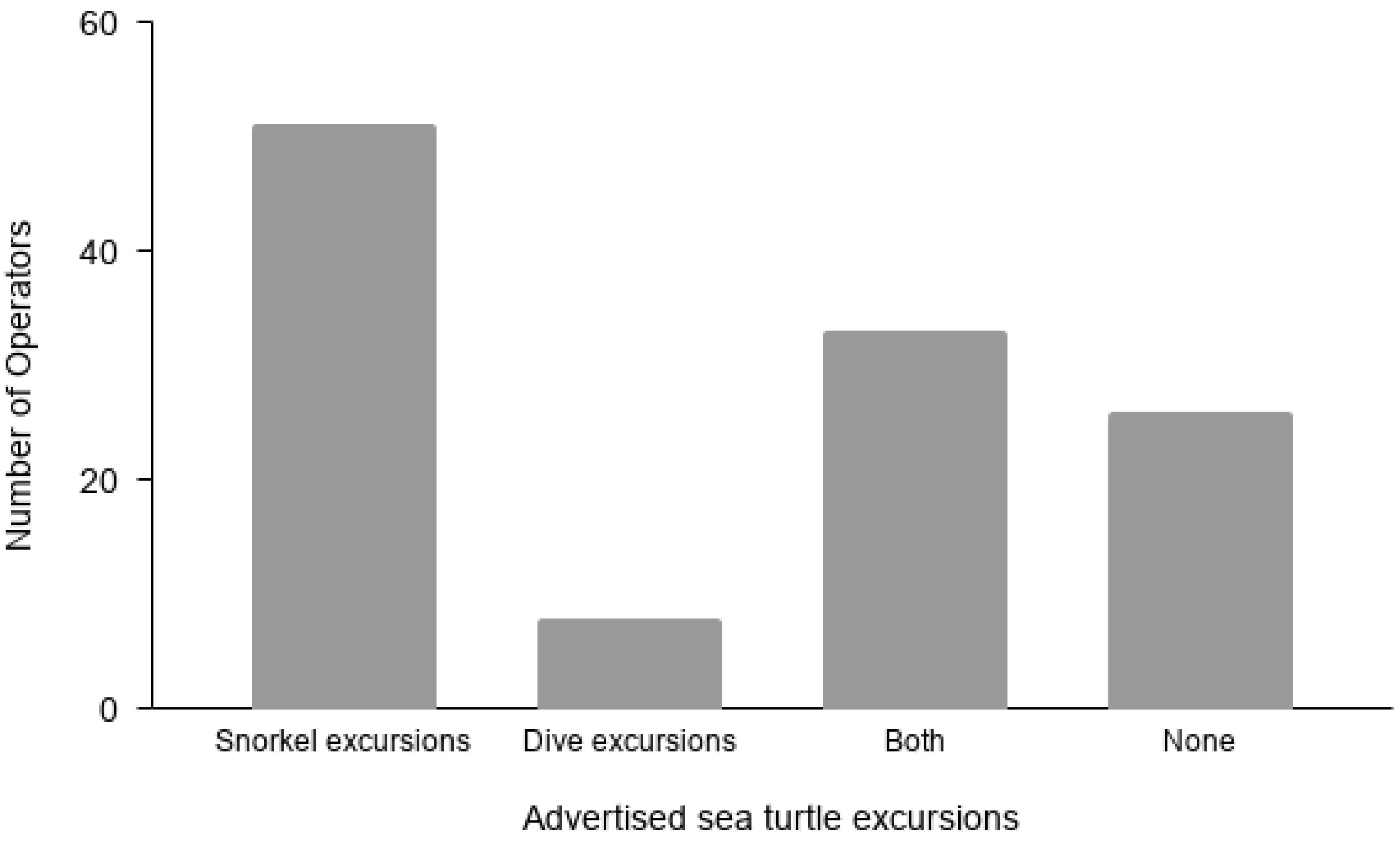
Number of operators that offered or did not offer advertised sea turtle excursions in 2022

### Sea Turtle Viewing

In 2022, an estimated 21,365 advertised sea turtle excursions were conducted, with snorkelling trips comprising 66.7% of all excursions, and dive trips making up the remainder. Snorkel excursions also accounted for 72% of the estimated 163,239 guests who participated in sea turtle trips that year.

### Value of Sea Turtle Tourism

In 2022, the direct revenue generated by sea turtle tourism in the Maldives was estimated at US$10.9 - 14.4 million. Sea turtle snorkel excursions on average generated substantially more revenue than dive excursions, accounting for 77% of the total direct revenue up to US$11.06 million, while sea turtle dives generated an estimated US$3.28 million. An estimated 797 jobs were directly supported by sea turtle tourism activities, 78.8% of which were held by Maldivian nationals (n=628).

Popular sites for sea turtle snorkeling (n=5) generated on average 250% more in estimated direct revenue than sea turtle diving sites (n=2), with snorkel sites such as Gemana Faru in Raa Atoll generating as much as US$605,384 in 2022 (Table 1). Comparatively, Kuredu Caves and Kuredu Express in Lhaviyani Atoll, the most popular sites for sea turtle dives, generated significantly lower revenues, averaging US$87,555 and US$82,680 respectively over the same period. The estimated number of trips and guests were also on average 141% and 46% higher respectively for snorkel excursions than dives excursions at these sites. Kuredu Caves is the only popular site where both snorkel and dive excursions are run, generating an accumulated US$383,254.

**Table 1.** Estimated economic contributions of sea turtle snorkeling and diving excursions across six popular sea turtle viewing sites (≥ 3 operator mentions) in 2022, categorized by the number of operators, trips, guests, and average revenue in US$.

| Site Name<br>(Atoll) | Snorkel Excursions |  |  |  | Dive Excursions |  |  |  |
| --- | --- | --- | --- | --- | --- | --- | --- | --- |
| | No. of Operators | Estimated No. of Trips | Estimated No. of Guests | Estimated Revenue (US\$) | No. of Operators | Estimated No. of Trips | Estimated No. of Guests | Estimated Revenue (US\$) |
| Biyadhoo Reef (Kaafu) | 3 | 572 | 4368 | 64,064 | - | - | - | - |
| Gemana Faru (Raa) | 5 | 832 | 6864 | 605,384 | - | - | - | - |
| Kuredu Caves (Lhaviyani) | 5 | 884 | 6708 | 295,698 | 4 | 416 | 4472 | 87,556 |
| Kuredu Express (Lhaviyani) | - | - | - | - | 3 | 364 | 4212 | 82,681 |
| Kuredu House Reef (Lhaviyani) | 6 | 1472 | 8175 | 332,538 | - | - | - | - |
| Makunudu Reef (Kaafu) | 6 | 936 | 5640 | 188,060 | - | - | - | - |

The maximum per turtle value at sea turtle viewing snorkel sites in 2022 was substantially higher than that for dive sites. At snorkeling sites, Gemana Faru had the highest estimated per turtle value at US$29,709, followed by Kuredu Caves, Makunudu Reef in Kaafu Atoll, and Kuredu House Reef in Lhaviyani Atoll (Table 2.). This was the highest observed across all popular sea turtle viewing sites. In contrast, the dive sites had lower estimated per turtle values, with Kuredu Caves and Kuredu Express in Lhaviyani Atoll, generating US$5,058 and US$4,733 per turtle, respectively. However, the combined value from snorkel and dive excursions at Kuredu Caves increases the per turtle value at this site to US$21,471.

**Table 2.** Estimated maximum economic value per turtle across six popular sea turtle viewing sites (≥ 3 operator mentions) in 2022. Only calculated for 75% of most sighted individuals at each site from the Olive Ridley Project sightings database for 2022.

| Site Name<br>(Atoll) | Snorkel Sites |  |  |  | Dive Sites |  |  |  |
| --- | --- | --- | --- | --- | --- | --- | --- | --- |
| | Estimated Revenue (US\$) | 75% Estimated Revenue (US\$) | No. of Individuals Sighted (75%) | Estimated per turtle value (US\$) | Estimated Revenue (US\$) | 75% Estimated Revenue (US\$) | No. of Individuals Sighted (75%) | Estimated per turtle value (US\$) |
| Biyadhoo Reef (Kaafu) | 72,800 | 54,600 | - | - | - | - | - | - |
| Gemana Faru (Raa) | 673,400 | 505,050 | 17 | 29,709 | - | - | - | - |
| Kuredu Caves (Lhaviyani) | 328,250 | 246,188 | 15 | 16,413 | 101,163 | 75,872 | 15 | 5,058 |
| Kuredu Express (Lhaviyani) | - | - | - | - | 94,663 | 70,997 | 15 | 4,733 |
| Kuredu House Reef (Lhaviyani) | 402,348 | 301,761 | 25 | 12,070 | - | - | - | - |
| Makunudu Reef (Kaafu) | 221,900 | 166,425 | 12 | 13,869 | - | - | - | - |

### Perceived Value of Sea Turtles

Sea turtles were frequently requested by tourists, with 92% of operators (n=107) reporting guest interest at least once per week, including 25% (n=29) who received daily requests. Only three operators reported never receiving requests to see sea turtles.

Operators ranked sea turtles higher for snorkelers than for divers among marine species that guests most wanted to see, with approximately 45% and 42% of snorkel operators ranking them as ‘Very High’ or ‘High’, respectively (Fig 3a). On the other hand, 81% of dive operators ranked sea turtles as either ‘High’ or ‘Medium’, listing other marine species such as manta rays (n = 45), whale sharks (n = 42) and other sharks (n = 17) as species divers wanted to see more than sea turtles (Fig 3b). Only 18% of operators ranked sea turtles as ‘Very High’ for divers, with 1 ranking them as ‘Low’. No operators ranked sea turtles as ‘Very Low’ for either snorkelers or divers.

**Figure 3.**
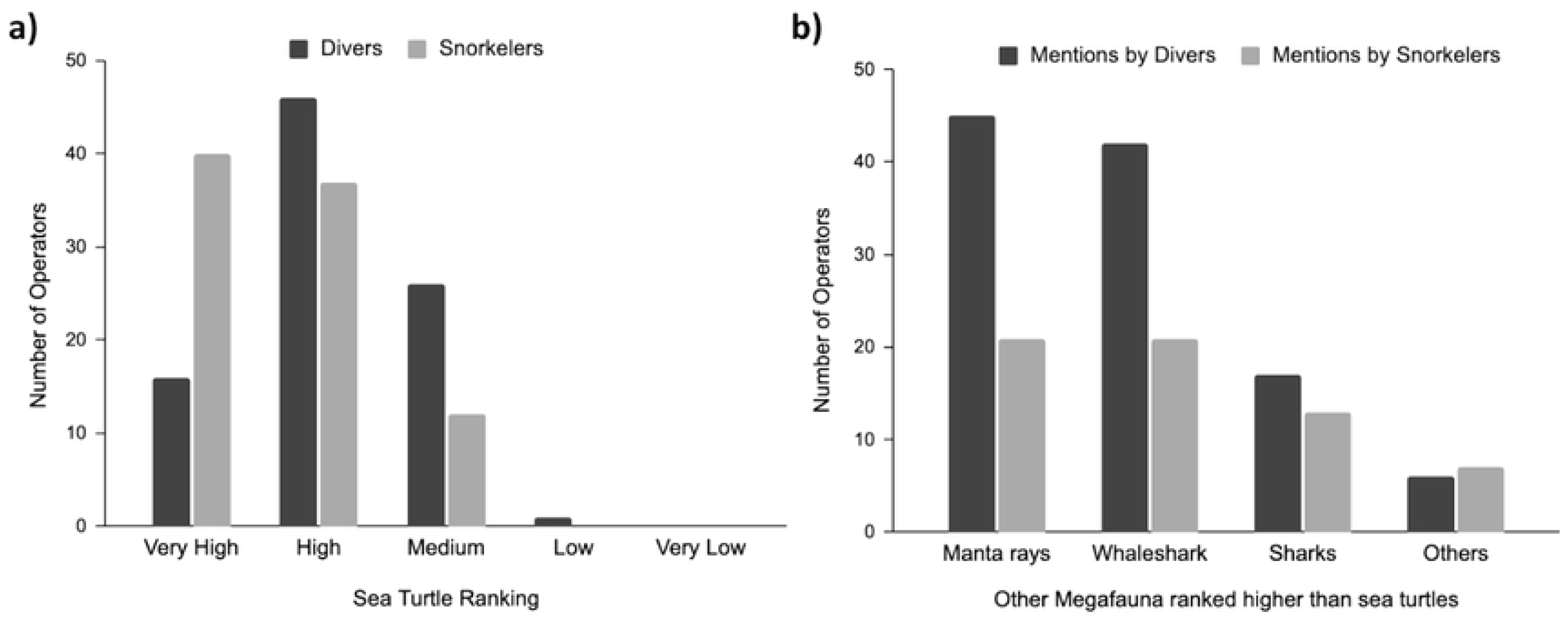
Survey responses on a) where sea turtles ranked among marine megafauna that guest most wanted to see and b) other megafauna that ranked higher than sea turtles, according to snorkel and dive operators

97.4% of operators (n = 113) stated that sea turtles were important for their business, and 110 operators featured sea turtles in their business operations for branding and marketing (Fig 4). The most common uses were ‘Media on Website’ and ‘Marketing Trips’, with over 70 operators featuring sea turtles through these channels. Sea turtles were also prominently featured in ’Art’ and ’Merchandise,’ with 35 and 27 operators incorporating them into these aspects of their business, respectively. Fewer operators featured sea turtles into physical structures or logos, with only six not using any sea turtle-related features in their operations.

**Figure 4.**
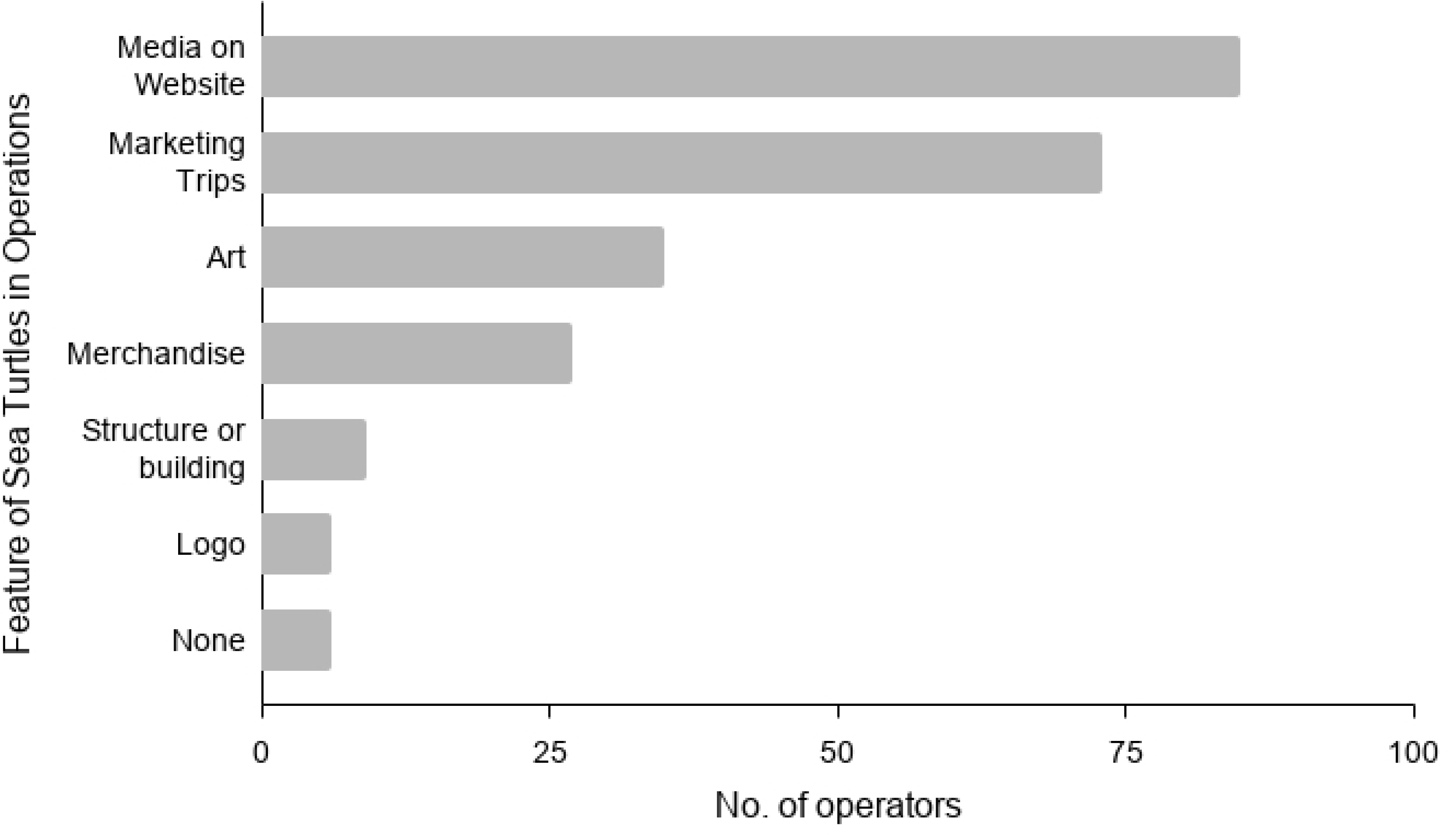
Feature of sea turtles in the operations of surveyed tour operators in 2022

Gemana Faru was listed by five operators as a frequently visited site for advertised sea turtle snorkelling excursions. Travel distances to this site ranged from 4 km to 12 km. For dive excursions, Kuredu Caves was listed by three operators, with travel distances ranging from 4 km to 22 km, the latter travelling from a neighbouring atoll (Fig 5).

**Figure 5.**
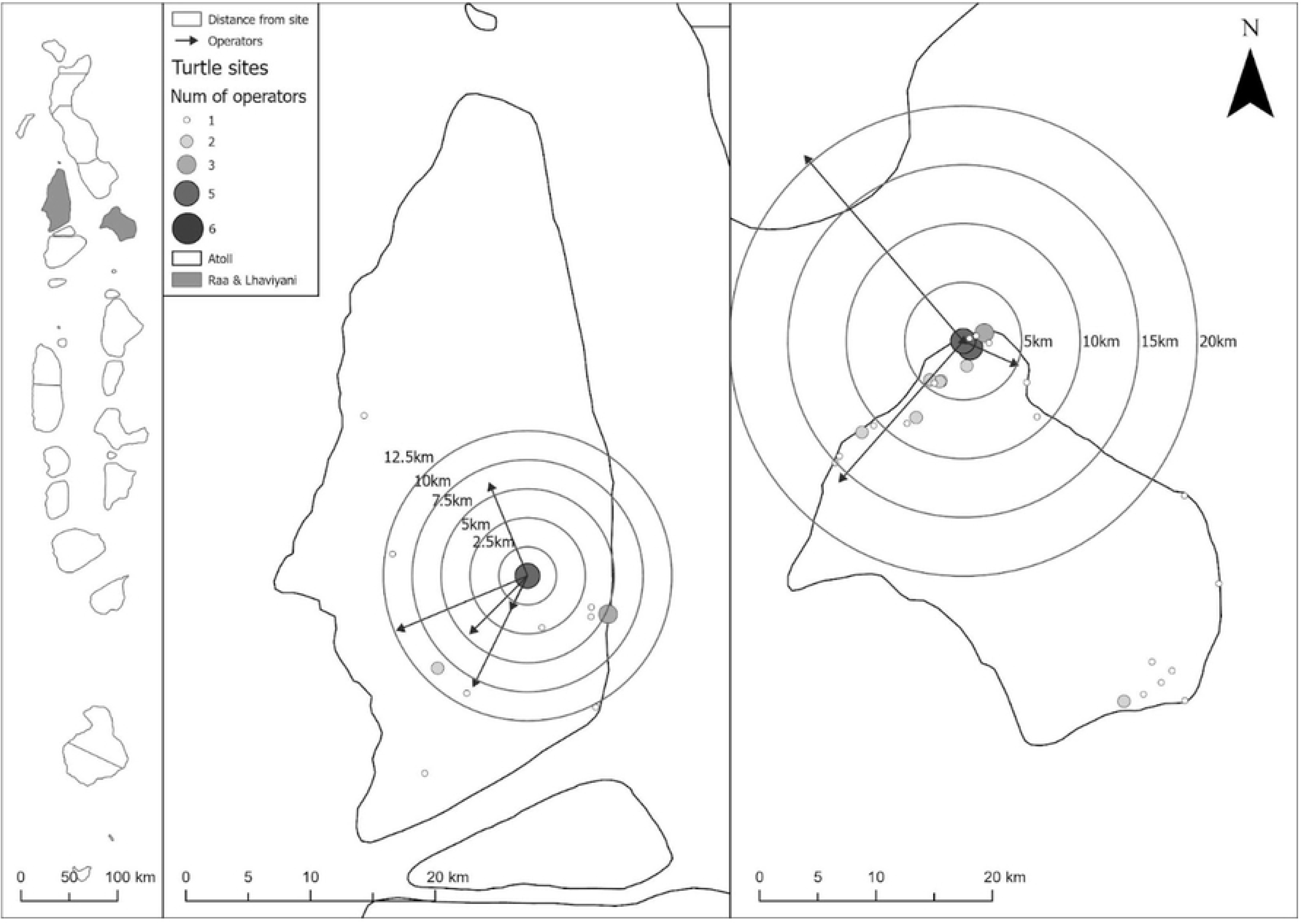
Map showing the distances traveled by tour operators to the most economically productive sites for sea turtle snorkelling (left; Gemana Faru, Raa Atoll, average income US$ 605,384) and diving (right; Kuredu Caves, Lhaviyani Atoll, average income US$295,698). The arrows and buffer rings indicate the travel distance for each operator. The Maldives/Atolls are Administrative atoll boundaries from planning.gov.mv accessed March 18th 2020.

## Discussion

### Value of Sea Turtle Tourism

The non-consumptive value of sea turtles is a major contributor to the Maldives tourism industry, generating up to US$14.4 million in estimated direct revenue in 2022, comparable to that of other charismatic marine megafauna such as manta rays [51] and sharks [52, 58]. This value is magnitudes higher than the US$1.08 million reported in 2019 [53] primarily due to a greater survey effort, producing a more comprehensive estimate of sea turtle tourism value, not only by the number of survey respondents, but also by geographic location. The significant non-consumptive value of sea turtles has been documented in other locations across the tropics [17–19, 40, 48, 59]. In Boa Vista Island, Cape Verde, the total economic value (TEV) of sea turtles was estimated at almost €2 million in 2019, with the non-consumptive value from ecotourism activities and conservation investment accounting for >99% of this value [47]. Similarly, a study by Teh et al. [49] estimated the TEV of sea turtles in Sabah, Malaysia to be US$23 million per year, almost all of which was associated with non-consumptive use through ecotourism. Consumptive uses (i.e. turtle egg consumption and sale) only accounted for 0.05% of this total value. Moreover, sea turtle tourism in Sabah supported an estimated 1146 jobs, equivalent to US$469,000 in employment income per year. In the Maldives, sea turtle tourism also contributed significantly to tourism employment, particularly for Maldivian nationals, who held 78.8% of the 797 jobs associated with this sector. These findings highlight sea turtle tourism as a major source of sustainable economic gain and employment, with strong potential to drive local support for conservation measures.

It is important to note that, although the response rates for this survey were marginally higher compared to the 2019 survey, the estimated value of sea turtle tourism was derived from less than half of operational resorts and only a small proportion of local island operators. As such, the true value of sea turtle tourism is likely considerably higher. Additionally, these estimates do not include the indirect revenue generated from sea turtle tourism (e.g. accommodation, food, and transportation expenditures) or the value of ecosystem services that sea turtles provide, such as the regulation of seagrass and benthic reef biodiversity through grazing [60] and the input of nutrients to beach ecosystems through nesting [61]. The tourism value from sea turtle nesting and hatching experiences were also not quantified in this study, which at present, occur opportunistically on resort islands that have sea turtles nesting and are typically not a charged activity. However, unofficially reports indicate that it is a major attraction for tourists and can influence their decision on where to stay. Future studies should therefore aim to incorporate WTP surveys to quantify the tourism value of nesting and hatching experiences for tourists, as demonstrated in other studies [18, 47, 57].

### Value of an Individual Turtle - Hotspot example

This study provides the first ever estimate for the non-consumptive value of an individual turtle in the Maldives. These values varied between popular sea turtle viewing sites and type of excursion, with the highest estimated value at approximately US$30,000 per year for a single hawksbill turtle at Gemana Faru, Raa Atoll. While previous studies have quantified the non-consumptive value of sea turtle populations in a specified area, few have provided estimates for individuals. In Sabah, Malaysia, the TEV of a single turtle was an estimated US$23,000 per year, with over 93% of this value associated with non-consumptive use through ecotourism [49]. This underscores the substantial economic value that sea turtles generate through non-consumptive use, often far exceeding their consumptive value.

Within the Maldives, hawksbill sea turtles are the most commonly observed species, and a large proportion of identified hawksbill turtles are juveniles, which is indicative of the region’s importance as a recruitment and developmental habitat for the species [62]. Juvenile hawksbills exhibit strong site fidelity to their foraging sites and typically remain within these developmental habitats for multiple years [63–66]. Studies in the Caribbean [46] and the Atlantic Ocean [67] have documented a minimum residency period of up to 3.8 yrs and 7.3 yrs, respectively. However, ORP’s Photo-ID program has recorded juvenile hawksbills foraging on the same reefs in the Maldives for as long as 14 yrs (ORP, unpublished data). The overall residency time might exceed this period as it is currently limited by the overall timeframe covered by the dataset, which was initiated in 2013, and includes sporadic historical records dating back to 1990. Continued monitoring might provide insight into longer residency times in the future. Similar residency patterns have been observed in other parts of the Indian Ocean, such as the Seychelles, where juvenile hawksbills remained in the same foraging grounds for up to 18 yrs [68], or the Chagos Archipelago, where turtles were observed for up to 24 yrs [69]. Therefore, under the assumption that the individuals at Gemana Faru display similar residency periods, the non-consumptive value of a single juvenile hawksbill could be as much as US$420,000 (over 14 yrs) or more.

Similarly, green turtles demonstrate high site fidelity to their foraging grounds across both juvenile and adult life stages [70–74]. Studies of Caribbean populations have reported minimum residency periods at foraging sites ranging from 2 to 17 yrs [73, 75, 76], while in Brazil, Colman et al. [77] reported similar long-term residence periods of up to 11.2 yrs. In the Indian Ocean, Sanchez et al. [68] found that green turtles in Aldabra, Seychelles, may spend >8 yrs residing in the same foraging grounds. In the Maldives, ORP have recorded residency periods for green turtles as long as 13 yrs at sites such as Kuredu Caves in Lhaviyani, a major hotspot for this species (ORP, unpublished data), suggesting that the non-consumptive value of a single individual may be equal to or exceed US$280,000 over their residency.

No prior peer-reviewed studies have assessed the consumptive value of sea turtles in the Maldives (i.e. consumption and sale of turtle meat, eggs and other byproducts), but anecdotal evidence and technical reports confirm that there still is a market for these products [78]. As such, a direct comparison cannot presently be made, but the potential conservation implications do highlight the importance of this topic for future studies.

### Implications for Tourism and Conservation

Similar to the 2019 survey [53], advertised sea turtle snorkels accounted for most of the total estimated direct revenue, trips, and guests in 2022. Sea turtles also ranked higher amongst snorkel operators compared to dive operators regarding marine megafauna that visitors most wanted to see. This is likely due to the higher accessibility and availability of sea turtles on typical snorkel trips, especially when compared to more elusive species such as manta rays or whalesharks. Green and hawksbill turtles can easily be encountered in reefs, lagoons and seagrass meadows across the Maldives [31], with a recent study suggesting that populations for both species are stable and/or increasing at many reefs in the Maldives [62]. In addition to this apparent preference for sea turtles, tourists regularly request to see sea turtles from the majority of tour operators indicating a clear demand for sea turtle tourism, which will only grow with the recovery of tourism since the COVID-19 pandemic.

This rising demand coincides with the expansion of guesthouses and local island tourism, which, despite only making 29% of survey responses in 2022, has become a rapidly growing sector of the Maldives tourism industry. Between 2019 and 2022, the number of operational guesthouses on local islands increased by 55.8% despite the disruption caused by COVID-19, offering visitors more affordable alternatives to the luxury resorts that have dominated the Maldives’ tourism market for decades [23]. The growth of guesthouse tourism has created direct economic opportunities for island communities by enabling them to engage more actively in an industry that has conventionally had limited linkages with local island economies [79, 80]. This has supported the growth of small tourism-related businesses on local islands, such as restaurants, handicrafts, and tour operators [80].

Together, the greater impact of sea turtle tourism on snorkel operators and the growth of guesthouse tourism highlights a significant potential to develop this sector further, particularly on local islands. Unlike scuba diving, snorkeling is more affordable and accessible for both tourists and prospective local guides, making it a relatively low-barrier and high potential avenue for developing sustainable ecotourism in the Maldives.

The geographic distribution of revenue generated by sea turtle tourism further illustrates this potential for growth. The highest levels of revenue generated from both snorkel and dive excursions were concentrated in the central and northern-central atolls where tourism activity was highest, while southern atolls had comparatively lower revenue gain (see S1 Fig for details). This further highlights the potential for expanding sea turtle tourism alongside broader tourism development in these regions, thereby creating a more inclusive distribution of economic benefits across the country.

Developing sea turtle tourism does not only provide opportunities for employment and economic gain, but may also contribute towards shifting perceptions of the value of sea turtles as an extractive to a non-consumptive resource. This shift has been documented in other locations where sea turtle tourism has been successfully implemented, including Costa Rica [46; 81], Brazil [40], and Trinidad [82]. However, it is important to recognise that while high economic value may be an incentive for change of use, it does not necessarily translate into support for conservation measures. The incorporation of educational initiatives and informative campaigns are also crucial to raise awareness of the ecological and intrinsic value of sea turtles alongside their economic value, which in turn, encourages local community buy-in for their protection [40].

### Sea Turtle Viewing Hotspots

The surveys identified sea turtle viewing ‘hotspots’ frequented by tour operators where the likelihood of encountering sea turtles is high, with some operators reporting encounter rates of over five turtles per trip at certain sites. These aggregation sites have the potential to generate significant revenue for multiple operators in the region, as demonstrated by sites such as Gemana Faru and Kuredu Caves. Operators were also willing to travel large distances to access these sites, further highlighting the strong demand for sea turtle viewing excursions and the importance of sea turtle aggregation sites such as these as a key attraction for tourists in the Maldives. Furthermore, operators identified numerous additional known sites with ‘*medium*’ to ‘*high*’ likelihoods of sea turtle encounters that were not used for advertised sea turtle viewing trips. For example, Makunudhoo in Kaafu Atoll, Dhonfanu Thila and Kihaa from Baa Atoll, and Madivaru and Rasdhoo Beyru in Alifu Alifu Atoll were frequently mentioned sites, among others. The extent of these additional sites reemphasises the potential for expanding the capacity of sea turtle tourism in the country, particularly in atolls with anecdotal reports of high densities of turtles but low tourism development, such as Gaafu Alifu and Dhaalu Atolls.

These findings can also help to map the extent of tourism pressure at hotspot sites used by multiple operators, and subsequently, help inform stronger protection measures in these areas. While sea turtles in the Maldives are typically accustomed to human presence to varying degrees [33], a rise in sea turtle ecotourism and the increased volume of human encounters it brings can have adverse effects on the natural behaviour and wellbeing of sea turtles [83–86]. It is therefore crucial to consider stricter wildlife interaction regulations alongside any plans for expanding sea turtle tourism in the country. Moreover, by mapping out the aggregation sites listed by tour operators, it is possible to identify potential key habitats for sea turtles, building on our current understanding of species and population distributions in the Maldives. This knowledge can contribute to more accurate population estimates and advise future legislation and management plans for sea turtles by highlighting both ecologically and economically significant sites for protection.

## Conclusion

Despite the disruption caused by the COVID-19 pandemic, the socio-economic value of sea turtles in the Maldives remains a significant one, comparable to other charismatic marine megafauna species. This study provides the first comprehensive estimate for the non-consumptive value of a single turtle in the Maldives, which might play a pivotal role in informing future management plans and shifting perceptions of sea turtle value from an extractive to a non-consumptive use one. The accessibility of sea turtle tourism to both tourists and tour operators also highlights a significant potential to develop this sector as a sustainable source of economic gain, particularly on local islands where the popularity of guesthouse tourism is growing. While this study associated economic value to direct revenue from sea turtle tourism, the indirect benefits and ecosystem service values remain unknown. Moreover, the tourism value of sea turtle nesting and hatching experiences was not assessed in this survey, despite being a major attraction for many tourists. Future studies should incorporate indirect revenue estimates, ecosystem service valuation, and WTP surveys to obtain a more comprehensive understanding of the non-consumptive value of sea turtles in the Maldives.

## Acknowledgements

The authors would like to thank all participants of the survey for their involvement and support, as well as all additional Olive Ridley Project staff who helped contact operators to collect survey responses. The study was conducted under permit from the Environmental Regulatory Authority, formerly Environmental Protection Agency Maldives (permit number: EPA/2023/PSR-T04).

## Supporting Information

### S1 Appendix. Survey questionnaire

**S1 Figure. Estimated annual revenue generated from sea turtle tourism across Maldivian atolls in 2022, shown as combined revenue snorkel and dive excursions (left), dive excursions (middle), and snorkel excursion revenue from both (right).** The Maldives/Atolls are Administrative atoll boundaries from planning.gov.mv accessed March 18th 2020.

## Notes

### Competing Interest Statement

The authors have declared no competing interest.

